# Human colorectal cancer: upregulation of the adaptor protein Rai in TILs leads to cell dysfunction by sustaining GSK-3 activation and PD-1 expression

**DOI:** 10.1101/2023.06.23.546212

**Authors:** Tommaso Montecchi, Giulia Nannini, De Tommaso Domiziana, Chiara Cassioli, Federica Coppola, Maria Novella Ringressi, Fabio Carraro, Antonella Naldini, Antonio Taddei, Giuseppe Marotta, Amedeo Amedei, Cosima Baldari, Cristina Ulivieri

## Abstract

**Background:** The immunosuppressive tumor microenvironment (TME) of colorectal cancer (CRC) is a major hurdle for immune checkpoint inhibitor-based therapies. Hence characterization of the signaling pathways driving T cell exhaustion within TME is a critical need for the discovery of novel therapeutic targets and the development of effective therapies. We previously showed that i) the adaptor protein Rai is a negative regulator of T cell receptor signaling and T helper 1 (Th1)/Th17 cell differentiation; and ii) Rai deficiency is implicated in the hyperactive phenotype of T cells in autoimmune diseases.

**Methods:** The expression level of Rai was measured by qRT-PCR in paired peripheral blood T cells and T cells infiltrating tumor tissue and the normal adjacent tissue in CRC patients. The impact of HIF-1α on Rai expression was evaluated in T cells exposed to hypoxia and by performing chromatin immunoprecipitation assays and RNA interference assays. The mechanism by which upregulation of Rai in T cells promotes T cell exhaustion were evaluated by flow cytometric, qRT-PCR and western blot analyses.

**Results:** We show that Rai is a novel HIF-1α-responsive gene that is upregulated in tumor infiltrating lymphocytes of CRC patients compared to patient-matched circulating T cells. Rai upregulation in T cells promoted PD-1 expression and impaired antigen-dependent degranulation of CD8^+^ T cells by inhibiting phospho-inactivation of glycogen synthase kinase (GSK)-3, a central regulator of PD- 1 expression and T cell-mediated anti-tumor immunity.

**Conclusions:** Our data identify Rai as a hitherto unknown regulator of the TME-induced exhausted phenotype of human T cells.

**What is already known on this topic:**

- Tumor hypoxia contributes to establish an immunosuppressive microenvironment that unleashes T cell function and limits the efficacy of immunotherapy in CRC.
- The molecular mechanisms underlying the impact of hypoxia on the signaling pathways controlling PD-1 expression are unknown.

**What this study adds:**

- A Rai/Akt/GSK-3 axis regulates PD-1 expression following TCR/CD28 co-stimulation
- This study uncovers the signaling pathway controlling hypoxia-dependent T cell exhaustion.

**How this study might affect research, practice or policy:** Rai expression levels in TILs of CRC patients might be explored as a potential new biomarker of T cell exhaustion and a predictive biomarker for anti-PD-1 response.

## Introduction

Colorectal cancer (CRC) represents approximately 10% of all cancers and is a major cause of cancer-related death worldwide.^1^ Despite major advances in the treatment of CRC patients following the introduction of immune checkpoint inhibitor (ICI)-based immunotherapy, prognosis remains poor because of the limited effectiveness of ICI in these patients, who experience a high rate of metastasis.^1–3^ Notably, the clinical ICI benefits in CRC patients has been associated with a higher mutation burden and tumor neoantigen load which in turn promotes the elimination of tumoral cells by tumor-infiltrating T cells (TILs).^3–5^ In addition, accumulating evidence indicates a prominent role for the CRC microenvironment in promoting the acquisition of a severe exhausted stage by TILs that cannot be reverted by immunotherapy, thereby contrasting anti-tumor immunity.^6^

T cell exhaustion is a state of T cell dysfunction characterized by the progressive loss of effector functions as well as transcriptional and metabolic changes.^7^ Markers of T cell exhaustion include the increased surface expression of inhibitory receptors such as programmed death-1 (PD-1), cytotoxic T lymphocyte antigen-4 (CTLA-4), lymphocyte activation gene-3 (LAG-3), T cell immunoglobulin and mucin domain 3 (TIM-3), and T-cell immunoglobulin and immunoreceptor tyrosine-based inhibitory motif domain (TIGIT), which by binding to their ligands on antigen presenting cells or tumor cells inhibit T cell activation.^7^

To date, exhaustion has been ascribed to the persistent exposure of T cells to their cognate antigens, which in turn results in a prolonged activation of TCR signaling that leads to a sustained expression of inhibitory receptors.^7, 8^ However, some TME-associated factors, including nutrient deprivation, hypoxia and increased acidity have more recently emerged as main drivers of T cell exhaustion.^6, 9, 10^ Indeed, a key role for the hypoxia-inducible transcription factor 1 (HIF-1) in the induction of CTLA-4 and LAG-3 expression on tumor infiltrating CD8^+^ T cells,^11^ as well as of protein death ligand-1 (PD-L1) on cancer cells^12^ has been demonstrated. Accordingly, inhibition of HIF-1α or reduction of hypoxia has been reported to unleash T cell activity and sensitize cancer to ICI.^13, 14^ Despite this evidence implicating hypoxia in immune suppression and resistance to ICI therapy of solid tumors, the molecular mechanisms underlying the hypoxia impact on TIL function are largely unknown, including its impact on signaling pathways controlling PD-1 expression.^10, 13^

The serine threonine kinase glycogen synthase kinase (GSK)-3 has been identified as a key regulator of PD-1 expression in T cells and as a new promising target for anti-cancer treatment of solid tumors.^15, 16^ GSK-3 is constitutively active in resting T cells, where it blocks cell activation and proliferation.^17^ Following TCR and CD28 triggering it is inactivated by the PI3K/Akt-dependent phosphorylation of two serine residues, Ser21 in GSK-3α and Ser9 in GSK- 3β.^15,16^ Moreover, the inactivation of GSK-3 by small inhibitors or its deletion in mouse cancer models has been shown to enhance CD8^+^ T cell function, leading to tumor growth suppression.^16, 18^ The up-regulation of Tbet transcription, which in turn inhibits PD-1 and LAG-3 transcription, has been suggested as the molecular mechanism underlying the anti-tumor activity of GSK-3 inactivation or deletion in T cells.^19^ How TCR signaling and GSK-3 inactivation are connected with Tbet transcription and whether the hypoxic TME impacts on GSK-3 activity to promote TIL exhaustion remains to be determined.

We have previously demonstrated that Rai, a member of the Shc family of protein adaptors, is a negative regulator of T cell activation and proliferation by preventing the recruitment and activation of key TCR signaling mediators such as ZAP-70 and PI3K/Akt.^20, 21^ Moreover, by modulating early TCR signaling, Rai restrains T helper 1 (Th1)/Th17 cell differentiation,^22^ thus emerging as a crucial regulator of T cell function. Consistent with this function, Rai-deficient mice showed spontaneous activation of T cells and enhanced development of Th1 and Th17 cells.^21, 22^

Here we investigated whether Rai expression is altered in TILs of CRC patients and whether it participates in the signaling pathways driving T cell exhaustion. We found that Rai expression is increased in TILs in a hypoxia-dependent manner, and that Rai upregulation resulted in the maintenance of active GSK-3 following TCR and CD28 triggering, which in turn is responsible for the high levels of PD-1 on T cells.

## Methods

### CRC samples, healthy controls and cell lines

Eight patients with CRC (4 males and 4 females, mean age 83,5□years) were enrolled after obtaining informed consent and approval of the local ethical committee (Comitato Etico Area Vasta Centro). Cancer samples were classified as colorectal adenocarcinoma according to the TNM classification of colorectal tumors. All patients underwent surgical resection of the primary lesion but did not receive chemotherapy; patients with evidence of serious illness, immunodeficit, or autoimmune or infectious diseases were excluded. Biological and clinical characteristics of the patients are summarized in Table 1. Fresh surgical specimens of CRC tissue were dissociated in order to isolate TILs. Tissue pieces from each patient were obtained from two different sites, central tumour (CT) and adjacent healthy mucosa (HM). Tissue samples were dissociated with the Tumour Dissociation Kit, human (Miltenyi Biotech, UK) in combination with the gentleMACS™ Octo Dissociator (Miltenyi Biotech, GmbH) to obtain a gentle and rapid generation of single-cell suspensions. In parallel, heparinised venous blood samples were collected and peripheral blood (PBMC) samples were isolated by density gradient centrifugation. Then, lymphocytes were magnetically isolated from dissociated CT, HM and PBMC samples with anti-human CD3 microbeads (Miltenyi Biotech, UK) using an AutoMACS Pro Separator device (Miltenyi Biotech, GmbH).

**Table 1.** Clinical characteristics of colorectal cancer (CRC) patients.

| Patient ID | Sex | Age | Diagnosis | TNM staging |
| --- | --- | --- | --- | --- |
| **CRC1** | F | 82 | Adenocarcinoma | pT2N0 |
| **CRC2** | M | 83 | Adenocarcinoma | pT3N2bM1 |
| **CRC3** | M | 90 | Adenocarcinoma | T4bN0M1b |
| **CRC4** | F | 85 | Adenocarcinoma | T4bN0M0 |
| **CRC5** | F | 83 | Adenocarcinoma | T3N0 |
| **CRC6** | M | 84 | Adenocarcinoma | pT3N2b |
| **CRC7** | M | 83 | Adenocarcinoma | pT0 |
| **CRC8** | F | 78 | Adenocarcinoma | pT3N0 |

Healthy control T cells were purified from buffy coats collected at the Siena University Hospital after receiving informed consent and approval of the local ethical committee (University of Siena Ethical Committee Board). T cells were purified by negative selection using RosetteSep^TM^ Human T cell enrichment cocktail (STEMCELL Technologies) following manufacturer’s instructions, while PBMCs were collected by direct stratification of the whole blood sample using Lympholyte® Cell Separation Media (Cedarlane). The T lymphoma-derived Jurkat cell line, Jurkat T cell transfectants expressing Rai,^20^ and the Burkitt lymphoma-derived Raji B cell line were also used. Cells were counted using Trypan Blue and Neubauer chamber (cell viability 90%). Cells were cultured at 37°C, 5% CO_2_ in RPMI-1640-sodium bicarbonate medium (Merck), supplemented with 10% bovine calf serum (GE Healthcare HyClone) and penicillin 50 IU/ml.

### T cell nucleofection and cell treatments

5×10^6^ freshly isolated T cells were co-transfected with 0.5 μg of pMAX-GFP and 1 μg of either pcDNA3.1-Rai or empty plasmid using homemade buffer 1M (5 mM KCl, 15 mM MgCl_2_, 120 mM Na_2_HPO_4_/NaH_2_PO_4_ pH 7.2, and 50 mM mannitol)^23^ and program V-024 of an Amaxa Nucleofector II (Lonza). T cells were allowed to recover in complete RPMI-1640-sodium bicarbonate medium and transfection efficiency was evaluated after 24h by flowcytometry and/or western blotting. Transfected cells were then stimulated for 24h at 37°C with plate bound 1μg/ml of anti-CD3 (clone UCHT1, BioLegend #300402) and 2μg/ml of anti-CD28 (clone CD28.2, BioLegend #302902) for qRT-PCR analysis and for FACS analysis of PD-1 expression. The anti-CD3 + anti-CD28 coating was prepared in PBS, added to 12 or 24 well plates (Sarstedt) and then incubated for 5h at 37°C. After incubation coating was removed by PBS washing before cells were added. For FACS analysis of GSK-3 α/β and Akt phosphorylation T cells were stimulated for 5 minutes at 37°C in serum-free RPMI-1640-sodium bicarbonate with anti-CD3 (1μg/ml) and anti-CD28 (2μg/ml) antibodies. Jurkat transfectants were activated for 5 minutes at 37°C in serum-free RPMI-1640-sodium bicarbonate by crosslinking the anti-CD3 antibody (1μg/ml, clone UCHT1, BioLegend #300402) with secondary anti-mouse antibodies for immunoblot analysis of GSK-3 α/β and Akt phosphorylation.

### Hypoxic treatment

Human primary T cells were cultured for 24 hours in a hypoxia workstation INVIVO_2_ 400 (Ruskinn Technology Ltd), providing a customized and stable humidified environment through electronic control of CO_2_ (5%), O_2_ (hypoxia: 2%, ∼14□mmHg) and temperature (37□°C), as previously described.^24^ Normoxic controls were kept at atmospheric O_2_ levels (21% O_2_, 5%CO_2_ and 74%N_2_).

### Immunoblotting

Primary T cells (1×10^6^) and Jurkat cells (3×10^6^) were lysed in lysis buffer (20 mM Tris-HCl pH 8, 150 mM NaCl, 1% Triton X-100) in the presence of Protease Inhibitor Cocktail Set III (Cal BioChem) and 0.2 mg Na orthovanadate/mL. Proteins were resolved by SDS-PAGE and transferred to nitrocellulose membrane (GE Healtcare) for immunoblotting analysis. Immunoblots were performed using the following primary antibodies: anti-Rai (BD, #610643), anti-pGSK3α/β (Cell Signaling, #8566), anti-pAkt (Cell Signaling, #9271), anti-actin (clone C4, Millipore, #MAB1501) and peroxidase-labeled secondary antibodies (anti-rabbit cod: 111-035-144 and anti-mouse cod:115-035-146 Jackson ImmunoResearch Laboratories). Labeled Abs were detected using the ECL kit SuperSignal® West Pico Chemiluminescent Substrate (Thermo Scientific) and scanned immunoblots were quantified using the ImageJ software (version 1.43r).

### Flow cytometry

The surface expression of PD-1 was assessed by labeling transfected T cells with anti-PD-1 (Genetex) primary antibody and Alexa Fluor-647 anti-mouse secondary antibody (Thermo Fisher Scientific) following 24h of treatment with anti-CD3 and anti-CD28 antibodies as described in Primary T cell nucleofection and treatments section. Pospho-GSK-3α/β and pospho-Akt were quantitated following 5 minutes of stimulation in transfected T cells fixed and permeabilized with fixation and permeabilization buffers (Biolegend) and stained with anti-pospho-GSK-3α/β (R&D systems, #AF1590) or anti-pospho-Akt (Cell Signaling, #9271) primary antibodies and Alexa Fluor-647 anti-rabbit secondary antibody (Thermo Fisher Scientific). The frequency of CD8^+^ CD57^+^ population in freshly isolated T cells was assessed by using PE-conjugated anti-CD8 (clone RPA-T8, Biolegend) and PerCP-conjugated anti-CD57 (clone HNK-, Biolegend) antibodies. Samples were acquired on Guava Easy Cyte cytometer 6-2L base (Millipore) and analyzed with FlowJo software version 6.1.1 (TreeStar Inc.).

### Degranulation assay

For the degranulation assay freshly isolated T cells from healthy donors with a percentage of CD8^+^ CD57^+^ (effector T cells) population higher than 10% of total were used. Raji B cells were loaded with a mixture of SEA, SEB and SEE peptides (2μg/ml) (Toxin Technology) or 1% BSA for controls, and incubated for 1h at 37°C. Then, T cells transfected with pMAX-GFP and either pcDNA3.1-Rai or empty vector were added in a 10:1 ratio and incubated for 4 h in presence of APC anti-LAMP1 antibody (clone H4A3, Biolegend #328620). Monensin (2mM, Biolegend #420701) was added 1h after the incubation start to block vesicular recycling. At the end of the incubation cells were stained with PE anti-CD8 antibody (clone RPA-T8, Biolegend #301008) and the degranulation extent was measured among the GFP^+^ population as the percentage of CD8^+^ T cells expressing LAMP1 on their surface by flow cytometry analysis.

### RNA Purification and qRT-PCR

For all the experiments RNA was extracted from samples by using the RNeasy Plus Mini Kit (Qiagen) according to the manufacturer’s instructions, and RNA purity and concentration were measured using QIAxpert (Qiagen). Single-strand cDNAs were generated using the iScriptTM cDNA Synthesis Kit (Bio-Rad), and qPCR was performed using the SsoFastTM EvaGreen® supermix kit (Bio-Rad) and specific pairs of primers listed in Table 2. Samples were run in triplicate on 96-well optical PCR plates (Sarstedt AG). Values are expressed as ΔΔCt relative to housekeeping gene HPRT expression. For analysis of Rai transcript in PBLs of CRC and healthy controls the relative gene transcript abundance was determined using ΔCt method and normalized to HPRT.

**Table 2.**
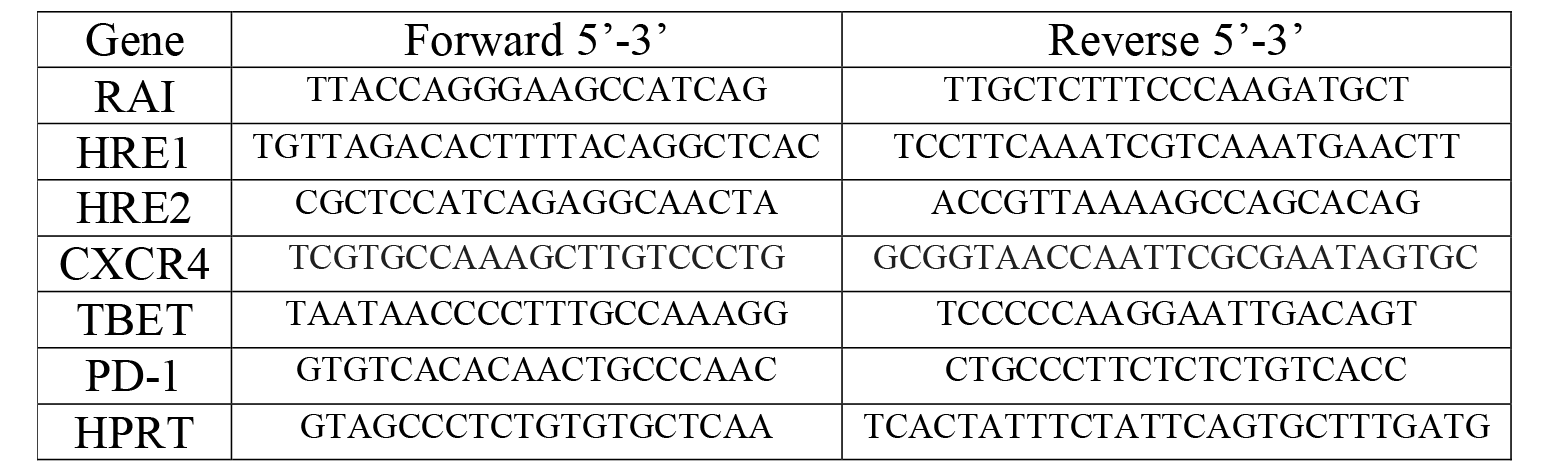
List of the primers used in this study.

### ChIP assay

Freshly isolated primary T cells (2×10^6^) from healthy donors were kept for 24 hours under hypoxic conditions and then were used to perform ChIP assay, according to MAGnify Chromatin Immunoprecipitation System protocol (Thermo Fisher Scientific), in order to assess HIF-1α binding to the two identified putative hypoxia responsive elements on *rai* promoter. In brief, cells were fixed with 1% formaldehyde for 10 minutes at room temperature then were lysed and exposed to 10 cycles of sonication (10 seconds on at 20% of power and 10 seconds off for each cycle) using SONOPULS Ultrasonic Homogenizers HD 2070 (BANDELIN) to obtain chromatin fragments of 200-500bp. Immunoprecipitations were performed using 5μg of anti-HIF-1α (clone PA3-16521, Invitrogen) or rabbit IgG as a control. The immunoprecipitated chromatin fragments were quantitated using qPCR. The primer pairs used to amplify HRE1 and HRE2 regions are listed in Table 2. Evaluation of the binding of HIF-1α to *cxcr4* promoter was used as a positive control of the assay.^25^

### RNA interference

Silencing experiments were carried out using a specific small interfering RNA (siRNA) targeting HIF-1α (Silencer®Select Validated siRNA, siRNA ID #s6541) and a control siRNA (Silencer®Select Negative Control #1siRNA Cat. n° 4390843), purchased from Ambion as previously described.^26^ Briefly, cells were seeded in a 6 well plate (Sarstedt) at a concentration of 2×10^6^ cells/well in RPMI 1640 without antibiotics, then transfection of 46 nM siRNAs was performed by using lipofectamine RNAi MAX (Invitrogen) diluted in OPTI-MEM® (1X) (Gibco, Thermo Fisher Scientific) according to the manufacturer’s instructions. Cells were incubated for 24h at 37°C under normoxic conditions (20.9% O_2_ and 5% CO_2_). Then the growth medium was replaced and supplemented with antibiotics, and cells were transferred for 24h to hypoxic conditions (2% O_2_ and 5% CO_2_) or maintained in a normoxic environment. Silencing efficiency was evaluated by Western Blotting, using HIF-1α primary antibody (BD Biosciences) and β-actin (Sigma-Aldrich). Anti-mouse IgG HRP (Cell signaling) was used as secondary antibody.

### Statistical analysis

Two-way ANOVA, Sidak’s multiple comparison was used for experiments in which multiple groups were compared. Wilcoxon matched-pairs signed rank test was used to compare matched samples and paired t test to determine the statistical significance of differences between two groups. GraphPad Prism Software (Version 8.4.2) was used for statistical analyses. A p < 0.05 was considered as statistically significant.

## Results

### Rai is differentially expressed in peripheral and TILs of CRC patients

We have previously demonstrated that Rai deficiency in mice is associated with T cell hyperactivation and autoimmunity.^21^ In addition, a decreased Rai expression was found in human peripheral blood lymphocytes (PBLs) from systemic lupus erythematosus (SLE)^22^ and multiple sclerosis (MS) patients (unpublished results), suggesting that inhibition of Rai expression in lymphocytes correlates with immune cell dysfunction in humans. Whether Rai expression is altered in T cells from cancer patients and associated with the reported exhausted phenotype is unknown.

To explore the hypothesis that Rai upregulation in T cells from cancer patients may contribute to the suppression of anti-tumour immunity we first compared Rai expression in PBLs from CRC patients and healthy controls by qRT-PCR. Rai was expressed at comparable levels between the two groups (figure 1A). Since T cells are reprogrammed by signals received from TME once infiltrated into the tumour mass we then asked whether the TME impacts on Rai expression in T cells. Quantification of Rai mRNA levels in CD3^+^ TILs from CRC patients and matched autologous PBLs revealed a significant upregulation in Rai expression in CD3^+^ TILs compared to PBLs (figure 1B). Notably, CD3^+^ TILs expressed higher Rai levels also compared with the CD3^+^ T cells infiltrating the matched normal tissue surrounding the tumoral mass (figure 1C). Additionally, analysis of Rai expression in CD3^+^ T cells in matched tumour, normal adjacent tissues and PBLs samples of a subset of patients further demonstrated that Rai was progressively upregulated in T cells moving from the periphery towards the tumour mass (figure 1D). Collectively, these data suggest that the TME positively impacts on Rai expression in T cells in human CRC.

**Figure 1.**
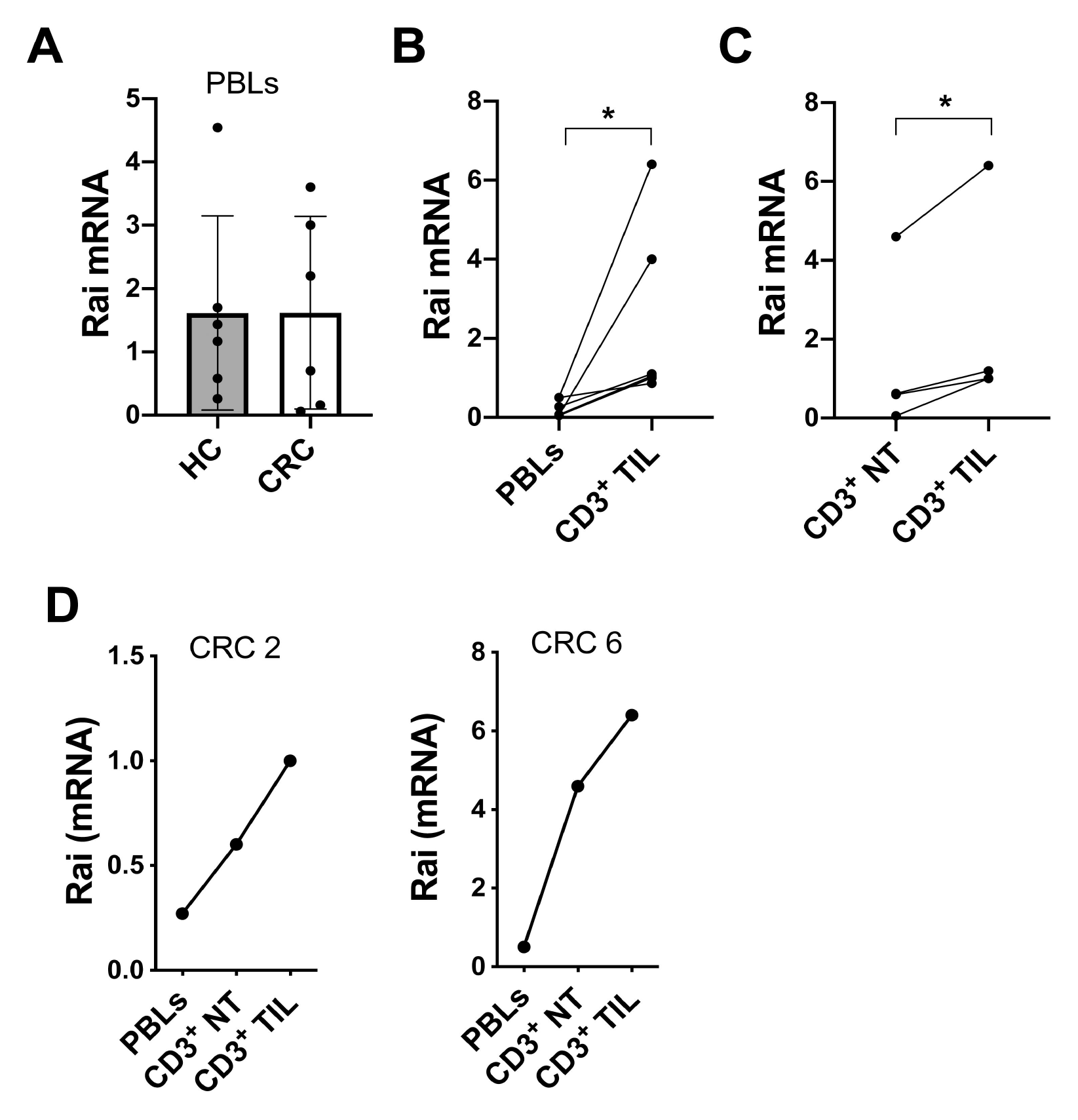
Expression of Rai in intratumor and normal adjacent tissue CD3^+^ T cells from CRC patients. (A) qRT-PCR analysis of Rai transcripts in PBLs of CRC patients (CRC) and healthy controls (HC). The relative gene transcript abundance was determined using dCt method and normalized to HPRT. (B, C) Comparison of Rai expression in PBLs, CD3^+^ tumour infiltrating lymphocytes (TIL) and CD3^+^ lymphocytes from tumor-adjacent normal tissue (NT) in matched samples from CRC patients. Data are presented as mean value ± SD (PBLs: n= 6 for CRC, n=7 for HC; TILs: n= 4, NT: n=4). (D) qRT-PCR analysis of Rai transcripts in PBLs, CD3^+^ lymphocytes in tumor-adjacent normal tissue (NT) and in TILs of 2 CRC patients. Wilcoxon matched-pairs signed rank test (B), paired t test (C). * p ≤ 0.02.

### Hypoxia-inducible factor-1**α** regulates Rai expression in T cells

Since hypoxia is a hallmark of solid tumors including CRC,^27^ where the partial oxygen pressure (pO_2_) drops in the tumour mass compared to normal adjacent tissues (median tumour pO_2_=19 mmHg in rectal carcinoma *vs* median normal tissue pO_2_=52 mmHg),^28^ and cell adaptations to hypoxia are primarily mediated by transcription factor HIF-1, we hypothesized that this transcription factor might account for the different Rai expression levels in PBLs and TILs of CRC patients. We thus evaluated Rai expression in CD3^+^ T cells from healthy controls (HC) subjected to hypoxia (2% O_2_ ∼ pO_2_14 mmHg). Hypoxic treatment significantly enhanced Rai levels, both as mRNA and protein, compared to normoxic controls (figure 2A).

**Figure 2.**
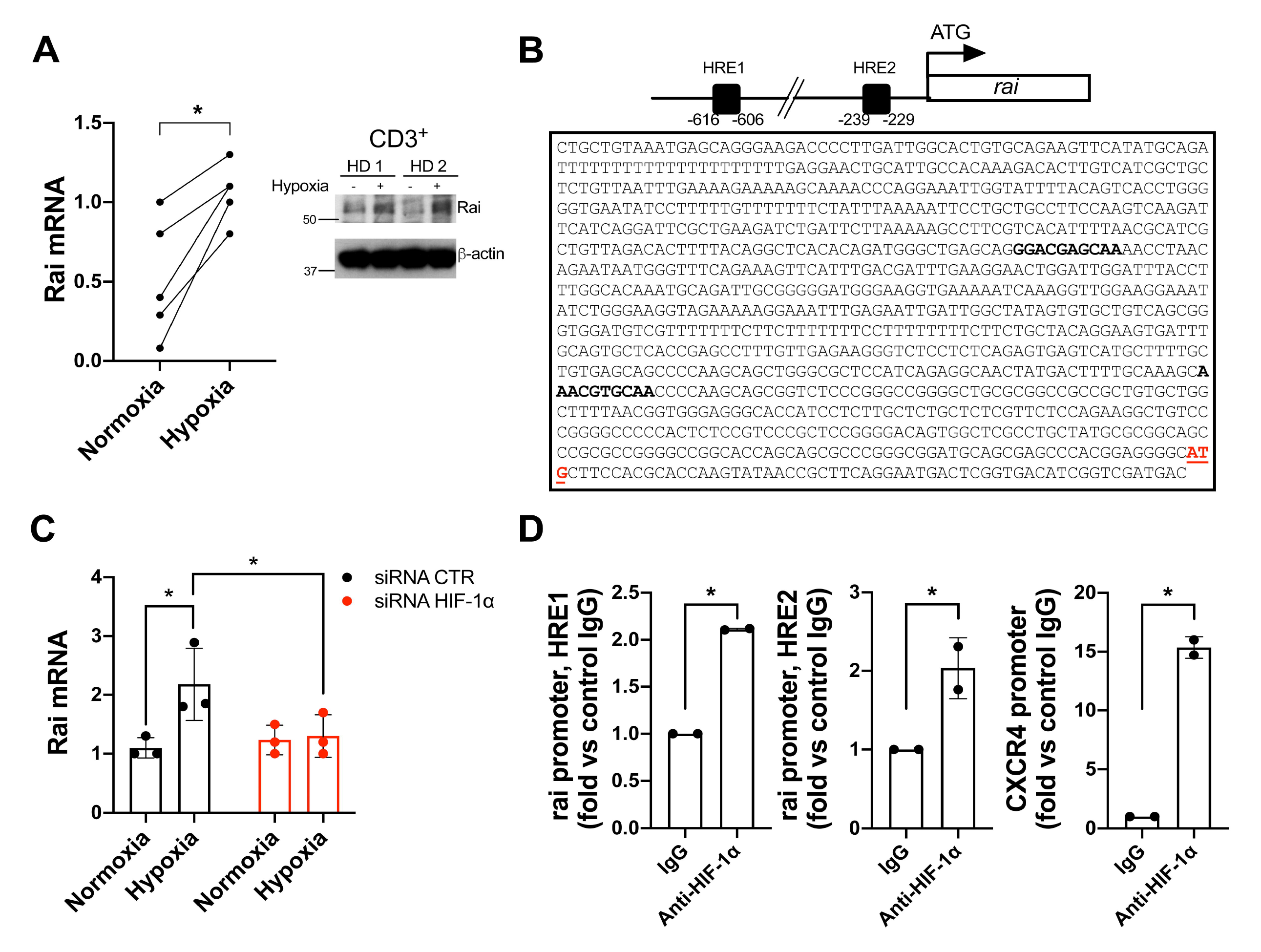
HIF-1α induces Rai expression in CD3^+^ T cells by directly interacting with the Rai gene promoter. (A) qRT-PCR and immunoblot analysis of Rai expression in CD3^+^ lymphocytes isolated from blood cultured under normoxic (Norm) or hypoxic (Hypo) conditions for 24 hr. Paired t test (qPCR: n = 5). * *p* = 0.01. (B) Schematic representation and sequence of the human *rai* promoter (nucleotides −958 to + 1). Black squares in the scheme and bold character in the sequence indicate the position of hypoxia-response elements (HRE) predicted by *in silico* analysis (score > 7). (C) qRT-PCR analysis of Rai transcripts in CD3^+^ lymphocytes silenced (siRNA HIF-1α) or not (siRNA CTR) for HIF-1α and cultured under normoxic (Normoxia) or hypoxic (Hypoxia) conditions for 24 hr. Data are presented as mean value ± SD (n = 3 healthy donors). Two-way ANOVA, Sidak’s multiple comparison test. * p ≤ 0.02. (D) ChIP assays of nuclear extracts of CD3^+^ lymphocytes cultured under hypoxic (Hypo) conditions for 24 hr using either anti-HIF1α or control unspecific Rabbit IgG (IgG) antibodies. Selected regions of *rai* promoters containing putative binding sites for HIF-1α showed in (A) were amplified by qPCR. Data are presented as a percentage of input DNA (n =3).

HIF-1 consists of a constitutively expressed HIF-1β subunit and a labile HIF-1α subunit which in the presence of O_2_ is hydroxylated by prolyl hydroxylase-domain protein 1-3, ubiquitinated by von Hippel-Lindau (VHL) proteins and degraded by the proteasome.^29^ Conversely, in hypoxic conditions, HIF-1α is not targeted for degradation and can translocate to the nucleus, where it dimerizes with HIF-1β and binds to hypoxia-responsive elements (HRE) on the promoters or enhancers of target genes, thus inducing their transcription.^29^

We performed an *in silico* analysis of the promoter region of the *rai* gene (0.9-kb, encompassing nucleotides −958 to + 1) to search for putative HIF-1α binding sequences using the JASPAR software. This analysis revealed two putative HREs on the *rai* promoter, namely HRE1 (from −616 to −606) and HRE2 (from −239 to −229), with a high relative profile score (> 80%) (figure 2B). Consistently, transfection of HC T cells with HIF-1α-specific siRNA significantly reduced Rai mRNA levels under hypoxia (figure 2C). To assess the direct binding of HIF-1α to the identified HRE regions in T cells, we performed chromatin immunoprecipitation (ChIP) assays using anti-HIF-1α antibody and the *cxcr4* promoter as a positive control.^25^ We found that HIF-1α binds to HRE1 and HRE2 under hypoxic condition (figure 2D), demonstrating that HIF-1α is recruited to the *rai* promoter and activates Rai transcription under hypoxia.

### Rai promotes PD-1 expression through the modulation of GSK-3α/β activity

The demonstration that Rai is upregulated in TILs of CRC patients, which are known to be dysfunctional and to express high levels of the immune checkpoint inhibitory receptor PD-1,^8^ together with the negative role played by Rai in antigen-dependent T cell activation,^20^ led us to investigate whether Rai impacts on the signalling pathways controlling PD-1 expression.

To dissect the role of Rai in this process and to mimic the hypoxia-dependent Rai upregulation, avoiding the global transcriptional changes induced by hypoxia, we overexpressed Rai in T cells. HC T cells were transiently co-transfected with a plasmid encoding Rai and a plasmid encoding GFP (figure 3A). Transfected cells were then stimulated by plate-bound anti-CD3 and anti-CD28 antibodies or left untreated, and the frequency of PD-1^+^ cells measured on GFP^+^ cells by flow cytometry. The frequency of PD-1^+^ cells was significantly increased in Rai-transfected cells compared to cells transfected with an empty vector following CD3/CD28 co-stimulation (figure 3B), suggesting a close relationship between increased Rai expression and PD-1 upregulation.

**Figure 3.**
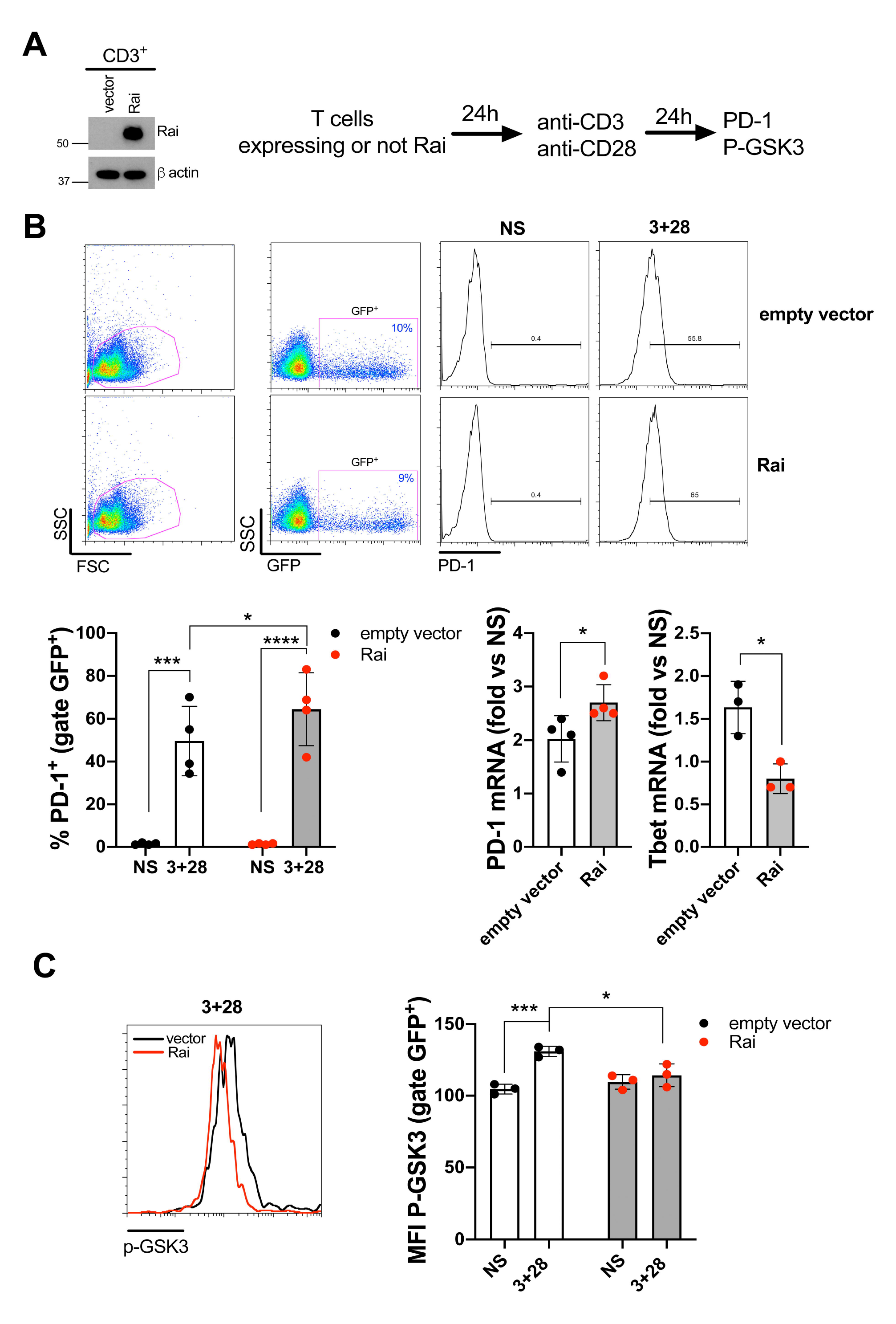
Rai promotes PD-1 expression by sustaining GSK-3α/β activity. (A) Representative immunoblot of exogenous Rai protein expression in CD3^+^ lymphocytes transiently co-transfected with the pcDNA3.1 and pmax-GFP plasmids (empty vector) or with the pcDNA3.1-Rai and pmax-GFP plasmids (Rai). (B) Flow cytometric analysis of the percentage of PD-1 positive cells (PD-1^+^) and qRT-PCR of PD-1 and Tbet transcripts in transfected T cells (GFP^+^) either untreated (NS) or activated with anti-CD3 and anti-CD28 (3+28) antibodies for 24 h. Representative flow cytometric histograms and gating strategy are shown. Data are presented as mean value ± SD. qRT-PCR data are presented as fold change of treated versus untreated sample (4 healthy donors, empty vector: 4 independent transfections, construct encoding Rai: 4 independent transfections). Flow cytometry data: Two-way ANOVA, Sidak’s multiple comparison test. * p < 0.05; *** p < 0.0002; **** p < 0.0001; qRT-PCR: unpaired t test. * p < 0.05. (C) Flow cytometric analysis of inactive GSK-3 (phosphorylated on Serine 9 and Serine 21, p-GSK3) in transfected T cells (GFP^+^) treated as in (B). Flow cytometric profiles of p-GSK-3 of Rai-transfected and mock-transfected cells following CD3+CD28 treatment. Data are presented as mean value ± SD (3 healthy donors, empty vector: 3 independent transfections, Rai: 3 independent transfections). Two-way ANOVA, Sidak’s multiple comparison test. * p < 0.05; *** p < 0.0007.

To define the mechanism linking Rai to PD-1 upregulation we focused on the enzyme GSK-3α/β, a central regulator of PD-1 expression in T cells, whose CD28/TCR-dependent phospho-inactivation has been shown to repress *pdcd1* transcription and to enhance cancer clearance.^15, 19^ Although a non-redundant role of GSK-3α and GSK-3β in T cells has been established using conditional knockout mice, with the β isoform playing a dominant role over the GSK-3α, inhibition of both isoforms has been recently shown to be required to reduce PD-1 expression and enhance T cell-mediated tumor rejection.^16^

We first examined by flow cytometry the phosphorylation status of the inhibitory residues on GSK- 3α/β in human primary T cell transfected with Rai-encoding or empty vector, following CD3/CD28 co-stimulation. Under these conditions, we observed a reduced basal phosphorylation of GSK-3α/β in Rai-overexpressing T cells compared with control (figure 3C). These results were confirmed by immunoblot analysis of Rai-expressing Jurkat transfectants (online supplemental figure 1). In addition, Rai expression also impaired TCR-dependent phosphorylation of both the α and β isoforms of GSK-3 and confirmed our previous data showing an impairment of Akt phosphorylation following TCR stimulation (online supplemental figure 1).^20^ In addition, qRT-analysis showed that Rai overexpression significantly increased PD-1 transcripts while reducing T bet transcripts, compared to control cells transfected with an empty vector (figure 3B). This finding is consistent with a report showing that enhanced GSK-3 phosphorylation promotes T bet upregulation which in turn decreases PD-1 expression in CD8^+^ T cells.^15^

Collectively, these results indicate that Rai promotes PD-1 upregulation by preventing CD3/CD28- dependent inactivation of GSK-3.

### Rai impairs antigen-dependent degranulation of CD8^+^ T cells

PD-1 is a marker of T cell exhaustion and plays a critical role in restraining tumour rejection by CD8^+^ T cells, which are endowed with the potential to recognize tumour antigens and to directly kill transformed cells trough the release of granules containing cytotoxic substances such as perforin, granzymes, tumour necrosis factor alpha (TNF-α) and membrane-anchored Fas-ligand.^30^

To determine the functional outcome of Rai overexpression in CD8^+^ T cells we evaluated the ability of these cells to degranulate in response to antigen stimulation, measured as surface expression of lysosome-associated membrane protein 1 (LAMP1) at the CD8^+^ T cell surface. CD3^+^ T cells, transfected with Rai-encoding or empty vector, were stimulated with Raji B cells loaded with bacterial superantigen (SE) and the frequency of CD8^+^ LAMP^+^ was quantified by flow cytometry within the GFP positive population, representing the transfected T cells. Rai overexpression led to a significant decrease of CD8^+^ degranulation when compared with control CD8^+^ T cells, showing that the ability of Rai to promote PD-1 upregulation translates into a functional defect (figure 4).

**Figure 4.**
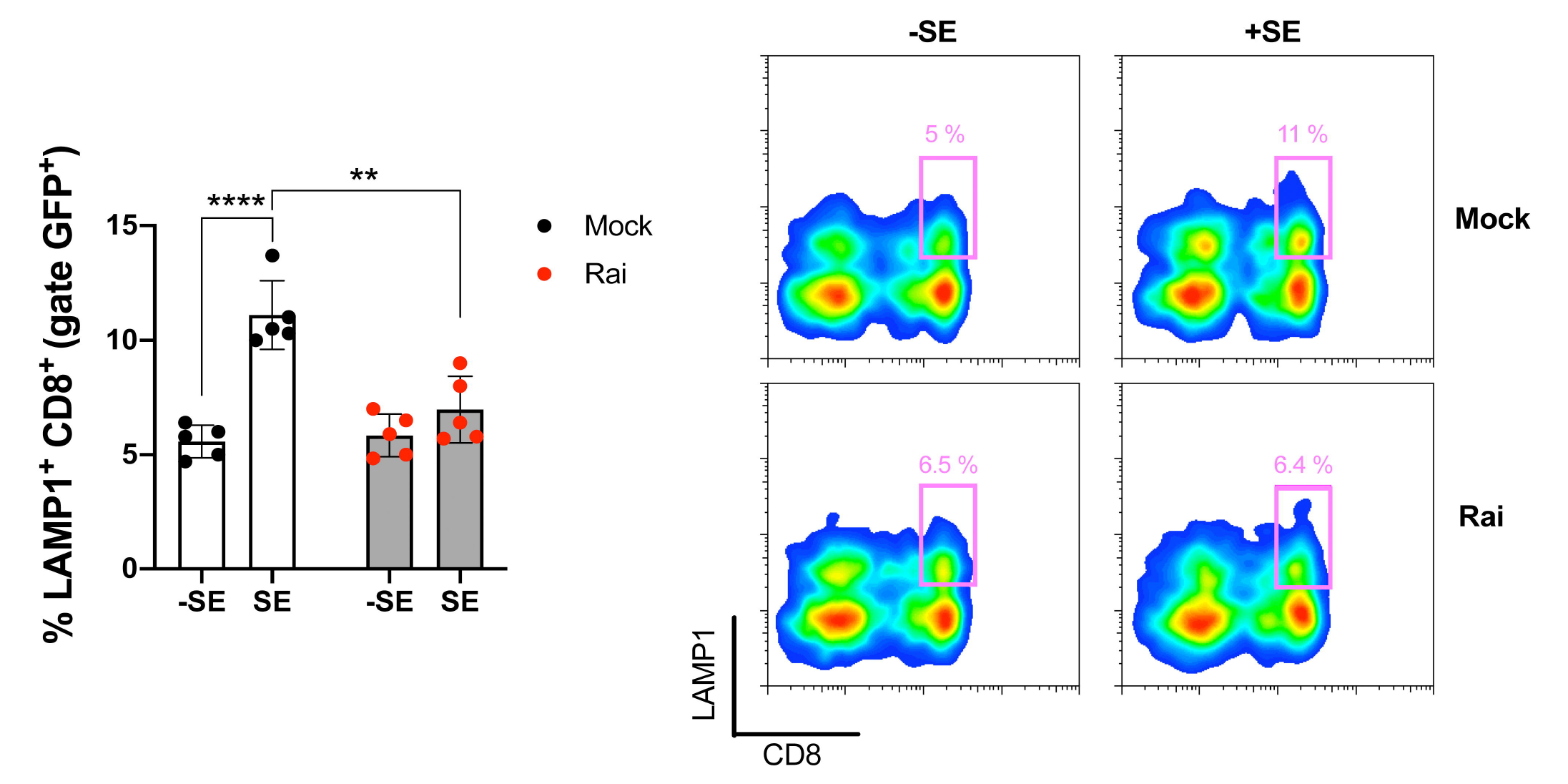
Rai overexpression leads to impaired degranulation of cytotoxic T lymphocytes expressing. (A) Quantification by flow cytometry of degranulation of CD8^+^ T cells among Rai-transfected and mock-transfected T cells (GFP^+^) incubated with Raji B cells loaded with a SAg mixture (+SE) or left unloaded (-SE). T cells were incubated with Raji cells at a 10:1 ratio. The histogram shows the percentage of LAMP1^+^ cells in CD8 T cells (LAMP1^+^CD8^+^) among transfected T cells (gated on GFP^+^). Representative flow cytometric dot plots with the CD8^+^LAMP1^+^ population delineated by the squares are also shown. Data are presented as mean value ± SD (5 healthy donors, mock: 5 independent transfections, Rai: 5 independent transfections). Two-way ANOVA, Sidak’s multiple comparison test. ** p < 0.002; **** p < 0.0001.

## Discussion

While the major role played by T cell-mediated immunity in CRC development^31–33^ suggests that this cancer could be targeted by immunotherapy, ICIs are ineffective in the majority of CRC patients because of the scarcity of immune cell infiltrates.^3^ Moreover, the TME may contribute to cancer immune escape and resistance to immunotherapy also in CRC subtypes with high levels of tumor immune infiltrates through several factors and processes including hypoxia, nutrient limitation, low pH and the presence of immunosuppressive metabolites and cells which, in addition to immune checkpoints, limit TIL functions.^6^ The complexity of the TME in CRC is a challenge for immunotherapy and the underlying mechanisms have only begun to be addressed.^3^ In this study, we identified a new mechanism that hinders the T cell anti-tumor response in the CRC tumor microenvironment.

A hypoxic TME is a major hurdle to the therapy of solid tumors. Indeed, a number of studies demonstrated that the enhanced tumor resistance to chemotherapy and aggressiveness relies on a mechanism involving the accumulation of the transcription factor HIF-1α in cancer cells where it promotes proliferation, migration and invasion. In addition, HIF-1α-dependent upregulation of the immunosuppressive ligand PD-L1 on tumor cells contributes to cancer immune escape.^34, 35^ Tumor immune escape is further favored by hypoxia-dependent release of lactic acid from tumor cells, which has been shown both to promote macrophage polarization towards the M2 subtype which secretes immunosuppressive cytokines and tumor-promoting soluble factors such as vascular endothelial growth factor (VEGF),^36^ and to suppress the cytotoxic activity of CD8^+^ T cells.^37^ In CRC, the importance of hypoxia-induced VEGF overexpression within the TME in driving chemoresistance is demonstrated by the finding that downregulation of HIF-1α expression overcomes the hypoxia-induced chemoresistance to oxaliplatin trough the reduction of VEGF levels and the inhibition of angiogenesis in a CRC mouse model.^38^ Moreover, a mechanism involving the VEGF-A-induced expression of the transcription factor TOX, a well-known marker of CD8^+^ T cell exhaustion,^39^ in TILs of CRC has been identified as one of the main causes of the resistance to anti PD-1 therapy, suggesting that reduction of VEGF-A levels in tumor tissue might also impact on T cell exhaustion.^40^ Consistent with this notion, the simultaneous blockade of PD-1 and VEGF-A in these patients restores antitumor function of T cells.^40^ HIF-1α inhibition has also been shown to abrogate PD-L1 mediated immune evasion by suppressing PD-L1 expression on tumor cells and tumor-associated macrophages, thereby fostering the implementation of combination therapies based on HIF inhibitors and anti-PD-L1 monoclonal antibodies.^41^

While the accumulation of lactic acid within TME has been suggested as the main mechanism underlying impaired T cell anti-tumor response driven by hypoxia, the impact of HIF-1α on T cell is as yet controversial. Opposite roles for HIF-1α in Treg and Th17 cell differentiation have been reported, with some studies demonstrating that HIF-1α inhibited Tregs and promoted Th17 cell differentiation and others showing, conversely, that HIF-1α promoted FoxP3 expression and Treg differentiation.^42, 43^ In addition, while stabilization of HIF-1α expression in murine CD8^+^ TILs through deletion of the von Hippel-Lindau gene has been shown to promote the expansion of a highly cytotoxic CD69^+^CD103^+^ Trm-like CD8^+^ TIL population and to improve the antitumor activity of CD8^+^ TILs in mouse models of solid tumors,^44^ other studies documented a direct role for HIF-1α in promoting expression of CTLA-4, LAG-3 and the loss of mitochondrial functions in T cells, leading to an impairment in their effector functions.^45, 46^ Interestingly, using breast cancer and colon adenocarcinoma mouse models, Bailey *et* al. recently demonstrated the efficacy of co-targeting HIF-1α and CTLA-4 in vivo, further highlighting a key role for HIF-1α in TILs.^12^ Although anti-PD-1 blockade, alone or in combination with chemotherapy or other ICIs, has been used in CRC, only some patients showed clinical advantage, and the mechanisms that underlying PD-1 resistance are still under investigation.^3^ In this context, while TME-dependent upregulation of PD-1 in TILs has been reported the role of hypoxia/ HIF-1α in this process has not been addressed.

Here we found that Hypoxia/HIF-1α, beside controlling PD-L1 expression on tumor cells by directly binding to the HRE motif in the promoter of PD-L1 gene,^12^ promotes PD-1 expression in T cells through a mechanism involving the upregulation of the cytosolic adaptor Rai, which our data identify as a novel HIF-1α target gene. In human T cells isolated from healthy controls the physiological levels of Rai were significantly increased in hypoxic condition compared with those under normoxia and conversely, when HIF-1α was knocked down, the effect of hypoxia on Rai expression was abrogated. Furthermore, we demonstrate the direct binding of HIF-1α to the *rai* promoter. Forced expression of Rai in human T cells led to enhanced PD-1 expression which correlated with reduced CD8^+^ T cell degranulation.

While previous work highlighted a key role of GSK-3 in PD-1 expression,^15, 19^ neither the upstream signaling events that govern TCR-dependent GSK-3 activity nor the impact of hypoxia on this process had been investigated. Our data identify a Rai/Akt/GSK-3 axis in PD-1 expression and Rai as a new regulator of GSK-3 activity under hypoxia in T cells.

The finding that Rai expression is significantly upregulated in TILs of CRC patients compared with matched circulating T cells, taken together with its impact on GSK-3 activity and PD-1 expression, further pinpoint Rai as a novel driver of T cell exhaustion within the TME. Therefore, HIF-1α inhibitors may represent an effective combination for PD-1-targeted immunotherapy. Based on our previous data showing that Rai overexpression in Jurkat T cell suppresses TCR signaling and T cell activation, together with the data presented in this report showing that upregulation of Rai in human T cells correlates with enhanced PD-1 expression and reduced CD8^+^ T cell effector function, we can speculate that different levels of Rai expression in TILs among CRC patients might contribute to the different response to anti-PD-1 therapy. Indeed, by directly inhibiting TCR signaling trough impairment of ZAP-70 activation^20^ and by regulating GSK-3 activity, Rai may prevent T cell reactivation within TME in the presence of anti-PD1 antibodies. In agreement with this hypothesis, melanoma patients sensitive to anti-PD-1 monotherapy showed a strong TCR signal strength signature.^47^ Moreover, while our data show a significant upregulation of Rai expression in TILs compared to PBLs or surrounding healthy tissue of the same patient, we documented a variability in Rai expression in TILs among CRC patients, suggesting that Rai expression levels in TILs of CRC patients might be explored as a potential new biomarker of T cell exhaustion and a predictive biomarker for anti-PD-1 response.

Although our study will need to be extended to a larger number of CRC patients, our data provide novel and encouraging insights into the hypoxia-dependent mechanism of T cell exhaustion in the TME and identify Rai as a novel regulator of this critical process in cancer development and treatment.

### Online Supplemental material

Included in the supplemental material is one supplementary figure showing the impact of Rai expression in CD3-dependent phosphorylation of AKT and GSK3 in Jurkat T cell.

### Author contributions

TM, GN, DD, CC, and FC did the experiments; TM, GN, FC, FCar, AN, AA, CTB and CU designed research and analyzed the data; TM and CU prepared the figures; TM, AA, CU and CTB wrote the paper. MNR, AT and GM provided essential reagents.

