## Supplementary figures and images for "Human colorectal cancer: upregulation of the adaptor protein Rai in TILs leads to cell dysfunction by sustaining GSK-3 activation and PD-1 expression"

### Supplemental figure 1

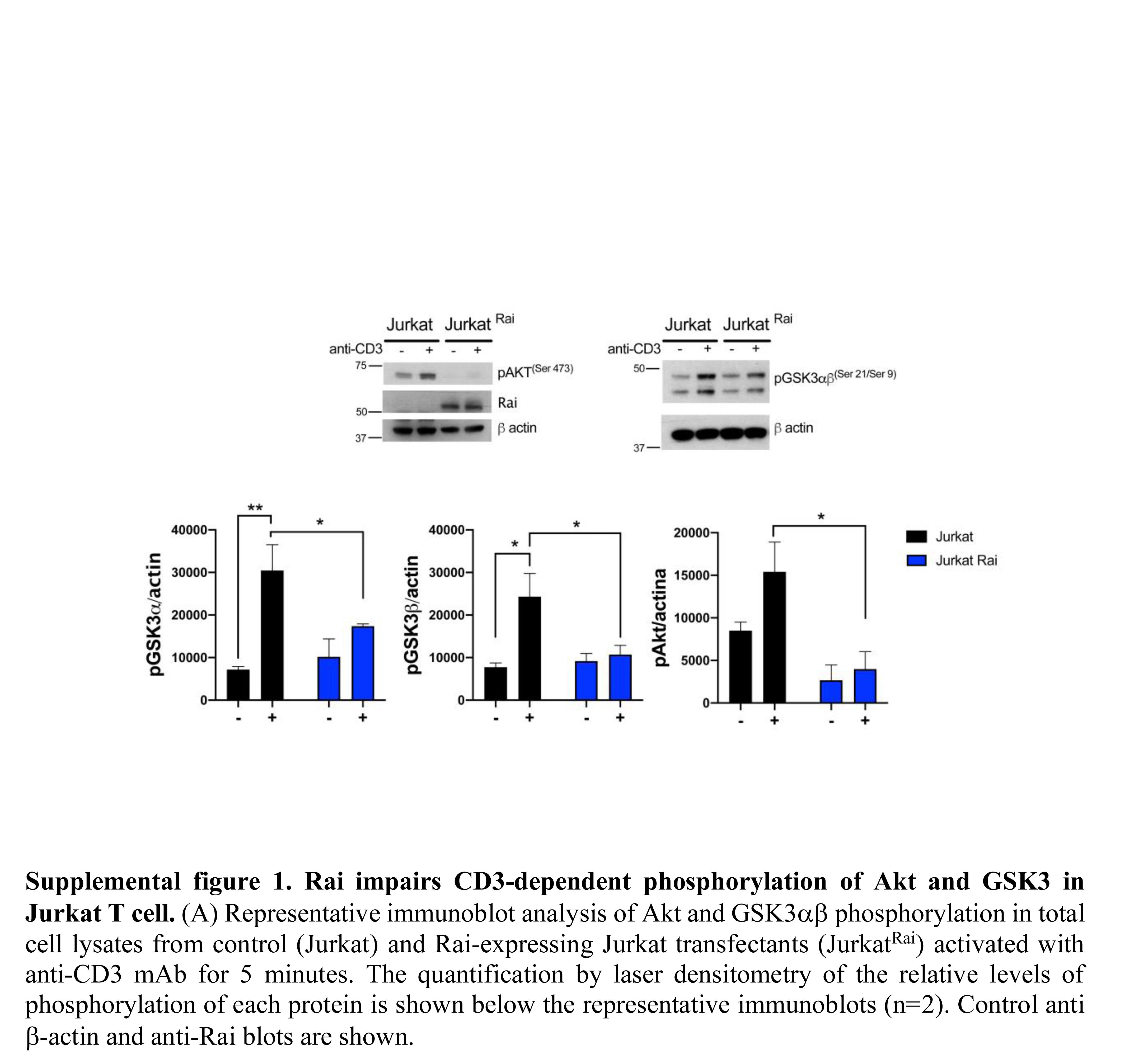
